# “Red cell enzyme polymorphisms in Muslim population of Eastern UP, India”

**DOI:** 10.1101/2022.11.24.517832

**Authors:** Pradeep Kumar, Vandana Rai

## Abstract

**Background:** Red cell enzyme polymorphisms have been used to study genetic variations in several human populations/countries worldwide. Owing to considerable ethnic and cultural heterogeneity in the Indian population, it is imperative to study different caste and religion of different regions.

**Aims and Objective:** To determine allele frequency of five Red Cell Enzymes (ADA, AK1, ESD, GLO1) in Muslim population of Eastern Uttar Pradesh

**Materials and methods:** Blood samples were collected from 200 unrelated individuals belonging to Muslim community of eastern Uttar Pradesh. The phenotypes of ADA, AK1, ESD and GLO1 systems were determined by agarose/starch gel electrophoresis. The allele frequencies were calculated by gene count method.

**Results:** The calculated frequencies of the alleles are as follows: ADA*1= 0.907, ADA*2= 0.092; AK*1= 0.92, AK*2= 0.08; ESD*1= 0.765, ESD*2= 0.235; GLO*1= 0.27,GLO*2 =0.73

**Conclusion:** The comparison of the allele frequencies of the four RBC enzymes studied in the present report with those of Asian populations showed that the allele frequencies are close to other Asian populations.

## Introduction

More than 50 different blood antigens are reported and few of them are very conveniently used for population genetic studies such as ABO. Since its discovery, frequency of ABO blood group is continuously studied in different global populations as well as in different religion and caste groups of India (Kumar et al., 2008,2009a,b,c,2010; Rai and Kumar 2010; Rai et al., 2009a,b,c,2010). Several publications reported association between ABO blood groups with different diseases such as malaria, COVID-19, cancer etc(Kumar and Rai,2012; Singh et al.,2019,2021). Similarly RBC blood cell enzymes are also polymorphic in nature and also convenient for population genetic studies, hence the present investigation aims to study four red cell enzymes (Adenosine deaminase, Adenylate kinase locus 1, Esterase D and Glyoxalase locus 1), whose polymorphisms are well established and standardized and well documented techniques are available. Variation in the distribution of phenotypes within different major ethnic groups of the world has been demonstrated for all the five enzymes studied.

Adenosine deaminase (ADA), catalyses the irreversible hydrolytic deamination of adenosine and deoxyadenosine to inosine and deoxyinosine. This enzyme is ubiquitous in mammalian and occurs in three molecular forms (Hirschhorn and Ratech, 1980). All three forms are coded by separate gene locus. Out of which, ADA1 (molecular mass of ~35 kDa) is a monomeric protein and its gene is located on chromosome 20 (20q 13.11) (Tischfield, 1974, Petersen et al., 1987; Jhanwar et al., 1989) and has two common alleles viz. ADA 1 (ADA*1) and ADA 2 (ADA*2). Spencer et al. (1968), first described the three genetically determined phenotypes (ADA1, ADA2-1 and ADA2). Fildes and Harris (1966) have observed polymorphism in AK1 isozyme and reported three distinct phenotypes: AK1, AK1, 2 and AK2. AK1 locus is present on chromosome 9(9q34.1- q34.3) having two alleles AK1*1 and AK1*2. Esterase D gene locus is assigned to chromosome 13 (13q 14.1-14.2) (Bender and Grzeschik, 1976), which is closely linked to retinoblastoma gene (Connolly et al., 1983). ESD locus contains two co-dominant alleles ESD*1 and ESD*2. The glyoxalase system consists of glyoxalase I and glyoxalase II enzymes. Glyoxalase I is dimeric protein present in the cytosol of cells. It catalyzes the formation of S-lactoylglutathione from the condensation of methylglyoxal and reduced glutathione glyoxalase II converts S- lactoylglutathione to D- lactic acid and reduced glutathione. In 1975, Kompf et al. (1975) reported polymorphism in GLO1 enzyme and observed three phenotypes: GLO1 1, GLO1 1, 2 and GlO1 2. The GLO1 gene is located on short arm of chromosome 6 (6p21.3-p21.2) (Bender and Grzeschik, 1976b) and locus is controlled by two co-dominant alleles GLO1*1 and GLO1*2.

Muslims of India make up more than 13.4% of the population (Census,2001). They belong to two major sects; Sunnis and Shias, while each sect has different Biradaris grouped under Ashraf and Ajlafs (Ansari,1959), these groups are based on traditional, social and occupational divisions. Ashraf comprises higher rank Muslims and Ajlaf includes lower rank Muslims like-Qureshi, Saifi, Ansari, and other lower occupation (Ahmad,1978). The aim of the present study is to investigate four red cell enzymes polymorphisms (ADA, AK1, ESD, GLO1) in eastern UP population belonging to a backward lower rank community of Sunni Muslims.

## Materials and methods

Ethical clearance was taken from Institutional ethics Committee of VBS Purvanchal University Jaunpur, India and informed written consent was taken from each subject prior to blood sample collection. 5ml blood sample was collected from 200 apparently healthy individuals selected from Muslim (Sunni) population inhabitants of UP. Blood samples were processed at the Human Molecular Genetics Laboratory, Department of Biotechnology, VBS Purvanchal University Jaunpur. Hemolysates were prepared by freezing and thawing method (Bhasin and Chahal, 1996). The samples were transferred into glass tubes (10×75mm) and centrifuged at 3000 rpm for 10 minutes to separate serum which was pipette out and the remainder (mainly red cells) was washed twice with isotonic (0.9% NaCl) saline to remove any traces of serum. Hemolysates were prepared by mixing an equal volume of distilled water to the washed packed red cells and mixing them mechanically using electric shakers. The mixture was kept overnight in a freezer (−20°C) where they were stored till their electrophoretic analysis. The isozymes of ADA and AK1 were electrophoresed on a single gel and stained simultaneously as described by Murch et al. (1986). The isozymes of ESD were electrophoresed following the technique of Wraxall and Storolow (1986). Mixed agarose/starch gel electrophoresis technique of Scott and Fowler (1982) was used for GLO1 typing.

## Results

The distribution of phenotypes and gene frequencies among Muslim population for four enzyme markers is presented in table 1 along with chi-square values for Hardy-Weinberg equilibrium. Total 200 Muslim samples are analyzed for ADA system. ADA*1 phenotype is found in 168 individuals, ADA*1,2 phenotype is found in 27 individuals and ADA*1,2 phenotype is found in 5 individuals. The overall phenotypic frequencies of ADA system are ADA*1 > ADA*1, 2 > ADA*2.The allelic frequencies of ADA*1 and ADA*2 are 0.9075 and 0.0925 respectively. In Muslim samples AK1*1 phenotype is observed in 171 samples, AK1*1,2 phenotype is observed in 26 samples and AK1*2 phenotype is observed in 3 samples. The overall phenotypic frequencies of ADA system are AK1*1 > AK1*1, 2 > AK1*2. The frequencies of AK1*1 and AK1*2 alleles are 0.92 and 0.08 respectively.

**Table 1.** Distribution of phenotypes of four enzyme systems (ADA, AK1, ESD, GLO1) in Muslim population

| Enzymes | Phenotype | Observed Number | Expected Number | Percentage | Allele Frequency |
| --- | --- | --- | --- | --- | --- |

|  |  |  |  |  |  |
| --- | --- | --- | --- | --- | --- |
| **ADA** | ADA*1 | 168 | 164.71 | 84 | ADA*1=0.9075 |
|  | ADA*1,2 | 27 | 33.58 | 13.5 | ADA*2=0.0925 |
|  | ADA*2 | 05 | 1.71 | 2.5 |  |
| **AK1** | AK1*1 | 171 | 169.28 | 85.5 | AK1*1=0.92 |
|  | AK1*1,2 | 26 | 29.44 | 13 | AK1*2=0.08 |
|  | AK1*2 | 03 | 1.28 | 1.5 |  |
| **ESD** | ESD*1 | 120 | 117.04 | 66 | ESD*1=0.765 |
|  | ESD*1,2 | 66 | 71.91 | 33 | ESD*2=0.235 |
|  | ESD*2 | 14 | 1.045 | 07 |  |
| **GLO1** | GLO81 | 16 | 14.58 | 08 | GLO*1=0.27 |
|  | GLO*1,2 | 76 | 78.84 | 38 | GLO*2=0.73 |
|  | GLO*2 | 108 | 106.58 | 54 |  |

ESD enzyme system is analyzed in 200 Muslim samples. ESD*1 phenotype is found in 120 samples, ESD*1, 2 phenotype is found in 66 samples and ESD*2 phenotype in 14 samples. The overall phenotypic frequencies of ESD system are ESD*1 > ESD*1, 2 > ESD*2. The allelic frequencies of ESD*1 and ESD*2 are 0.765 and 0.235 respectively. GLO enzyme system is analyzed in 200 Muslim samples. GLO*1 phenotype is present in 16 samples, GLO*1, 2 phenotype is present in 76 samples and GLO*2 phenotype is present in 108 samples. The percentage frequencies of phenotypes GLO*1, GLO*1, 2 and GLO*2 are 8%, 38% and 54% respectively. The frequencies of GLO*1 and GLO* 2 alleles are 0.27 and 0.73.

## Discussion

During the last five decades an impressive number of genetically controlled variations in red cell enzyme groups have been described and a fair amount of data is now available for genetic markers on different population groups of the world. Several studies were published for red cell enzyme marker polymorphisms in Muslim population from India but so far only one study was published from Uttar Pradesh.

In Muslim samples studied only ADA*1 and ADA*2 alleles were found no rare allele was observed in any samples. The frequency of ADA*1 was 0.907 and ADA*2 was 0.0925, which were comparable with the earlier reports of Lanchbury et al. (1996) about the Muslim population (ADA*1=0.909 and ADA*2= 0.0911). ADA*2 frequency as reported slightly lower in Shia Muslims (0.0767) in comparison to Sunni Muslims. The ADA*2 allele has shown a wide range of distribution in different Muslims population of India, varying from 0.091 in Uttar Pradesh to a surprisingly high value of 0.50 in Assam. Rare ADA*6 allele was reported among Muslims of Delhi. In earlier studies from other states the incidence of AK1*2 has been reported from a low of 0.062 in Sunni Muslims of Jammu and Kashmir to 0.120 in Muslims of Kerala. AK1*2 Allele frequency in Sunni and Shia Muslims reported 0.1177 and 0.1005 respectively.

The polymorphism of Esterase D enzyme has been studied in different populations of the world. In Muslim population inhabitant different states of India, the ESD*2 fequency has been reported in a range of 0.185 to 0.293. Chahal et al.(1989) reported GLO*1 allele frequency of 0.297 in Sunni Muslims from Jammu and Kashmir in Pulwama district whereas Bhasin et al (1992b) reported comparatively low frequency (0.205) from Srinagar Muslims. In a small samples of Delhi Muslims, Ghosh (1976) estimates the frequency of GLO*1 0.303 while Kumar and Rao (1982) reported higher frequency (0.327) from Hyderabad GLO*1 allele. In present study the frequency observed in the Muslims of eastern UP (0.27) fit well into the range reported by these studies.

## Acknowledgements

We are grateful to the subjects, who participated in this study without their cooperation this study could not be completed. We also gratefully acknowledge the financial assistance provided by University Grants Commission, New Delhi, India as major research project (Grant No. 39-968/2010(SR)) to Pradeep Kumar.

